# Real-Time Visualization of Calcium Phosphate Formation on Titanium Dioxide Nanoparticles Using Liquid Transmission Electron Microscopy

**DOI:** 10.1101/2025.04.16.648725

**Authors:** Jing Zhang, Liza-Anastasia DiCecco, Alyssa Williams, Alessandra Merlo, Kathryn Grandfield

## Abstract

The integration of titanium dioxide (TiO_2_) with calcium phosphate (CaP)-based hydroxyapatite (HAP) is a promising strategy for enhancing the bioactivity of bone implants. However, a fundamental understanding of the interfacial reactions governing CaP mineralization on TiO_2_ remains limited due to lack of characterization techniques with sufficient spatial and temporal resolution in hydrated state. In this study, we combined *in situ* liquid TEM imaging and correlative *ex situ* TEM analyses to investigate the nucleation, aggregation, and crystallization of the CaP layer on TiO_2_ nanoparticle surfaces. Our findings reveal a three-step mineralization process: (1) initial aggregation of TiO_2_ nanoparticles in solution, (2) formation of an amorphous calcium phosphate-like (ACP-like) layer on the TiO_2_ surface, leading to an ACP-like coated TiO_2_ nanoparticle structure, and (3) progressive crystallization of ACP into HAP, forming a HAP-like coated TiO_2_ nanoparticle structure. Liquid-TEM imaging captured dynamic transformations, including nanoparticle aggregation, structural evolution, and phase transitions, providing unprecedented insights into the physicochemical interactions underlying mineralization. Additionally, we evaluated the effects of electron beam exposure on TiO_2_ nanoparticles, demonstrating that high electron flux densities can induce morphological instability. This study advances our understanding of CaP-TiO_2_ interfacial mineralization and offers valuable guidance for optimizing bioactive coatings to improve osseointegration and the long-term stability of bone implants.

## Introduction

Bone implants are essential in modern orthopedic and dental medicine, serving to provide structural support, replace lost bone tissue, and facilitate bone regeneration. Metallic implants, particularly those composed of titanium (Ti) and its alloys, are widely utilized due to their exceptional mechanical strength and resistance to degradation in physiological environments.^1,2^, However, a major limitation of Ti-based implants is their bio-inert nature, which prevents direct structural and functional integration with surrounding bone tissue—an essential requirement for successful osseointegration.^3,4^

The naturally occurring or synthetically augmented Ti dioxide (TiO_2_) layer on Ti implants acts as a bioactive interface, facilitating cell attachment and bone matrix deposition.^5^ Recent advances in nanostructured TiO_2_ surfaces, such as nanotube or nanoporous coatings, have further enhanced protein adsorption and cell interactions, leading to accelerated bone healing.^6–10^ TiO_2_ also plays a crucial role in promoting calcium phosphate (CaP)-based hydroxyapatite (HAP, Ca₁₀(PO₄)₆(OH)_2_) formation,^11–13^ the primary inorganic mineral that makes up 65% of all human bone.^14^ HAP-coated TiO_2_ surfaces on Ti implants aim to enhance both mechanical stability and bioactivity by integrating the favourable mechanical properties of Ti and the biological properties of calcium phosphate ceramics.^15–17^ In an *in vivo* study, researchers implanted HAP–gelatin nanocomposites integrated with macroporous TiO_2_ scaffolds in rats.^17^ The results demonstrated that at 8 and 12 weeks post-implantation, the engineered scaffolds exhibited superior mechanical strength and enhanced osteointegration compared to natural calvaria bone.^17^ A clear understanding of the interfacial reactions between CaP and TiO_2_ is crucial for advancing knowledge on bone-implant interactions at the fundamental level, paving the way for developing next-generation biomaterials with optimized bioactivity and mechanical performance. However, direct, real-time validation of these reactions remains limited, as most studies rely on indirect or *ex situ* methods,^18–20^ leaving key mechanistic aspects debated.

*In situ* liquid transmission electron microscopy (TEM) has emerged as a powerful technique for examining biological materials in their hydrated, near-native state.^21^ Unlike traditional or cryogenic TEM methods, which provide only time-stamped static images of dry or frozen samples,^22^ liquid-TEM enables real-time observation of dynamic processes such as precipitation,^23^ particle movements,^24^ and surface disposition process.^25^ This novel approach has facilitated breakthroughs in the visualization of various biological structures,^211^ such as soluble proteins aggregate^26^, whole cells^27–29^ and the formation of biologically relevant minerals.^30,31^ In the field of bone implant research, liquid-TEM offers a unique opportunity to study real-time reactions at the bone-implant interface at the nanoscale,^32,33^ for example, our recent work has combined focused ion beam (FIB) nanofabrication of Ti lamellae with *in situ* liquid-TEM, allowing us to visualize CaP-Ti interfacial mineralization events directly on titanium.^32,33^

A common method for preparing liquid samples for TEM involves encapsulating the liquid specimen between electron-transparent silicon nitride (SiNx) windows, which are generally up to 50 nm.^34,35^ While this setup enables imaging under liquid conditions, a key challenge is controlling the liquid thickness to minimize electron scattering. Because the electron beam must pass through both SiNx membranes and a significant liquid thickness before reaching the detector, sample dimensions must be at the nanometer scale.

Here, we selected TiO_2_ nanoparticles (NPs) as a system to model the naturally occurring TiO_2_ oxide layer or synthetic coatings applied to biomaterials surfaces and employed liquid-TEM to directly observe and analyze interfacial reactions between CaP and TiO_2_ in real-time. This advanced imaging technique enabled direct visualization of ACP-coated TiO_2_ nanoparticle aggregation and its subsequent crystallization into HAP on the TiO_2_ surface, capturing dynamic transformations previously inaccessible through conventional characterization techniques. Our findings provide deeper insights into the physicochemical interactions governing bone-implant integration and offer valuable guidance for optimizing bioactive implant coatings to enhance osseointegration and long-term implant stability.

## Experimental Methods

### Calcium phosphate mineralization solution preparation

The calcium phosphate (CaP) mineralization solution was prepared based on a modified version of the method reported by Deshpande and Beniash.^36^ To achieve the desired final concentrations, equal volumes of three separate solutions were combined: 6.8 mM CaCl_2_·2H_2_O in 50 mM Tris (pH 7.4, 37°C), 36 mM Na_2_HPO₄ in 50 mM Tris (pH 7.4, 37°C), and 100 mM Tris–500 mM NaCl (Tris-NaCl, pH 7.4, 37°C). Subsequently, di-water or TiO_2_ nanoparticle suspension was added, resulting in a final solution containing 1.7 mM CaCl_2_·2H_2_O, 9 mM K_2_HPO₄, 50 mM Tris, and 125 mM NaCl. Prior to use, all reagents were passed through a 0.2 μm acrodisc syringe filter to reduce large aggregates.

### Titanium (IV) oxide (TiO_2_) nanoparticle suspension preparation and mineralization

TiO_2_ NPs (<25 nm, product number 637254) were sourced from Sigma Aldrich, Canada. A TiO_2_ nanoparticle suspension was prepared at a concentration of 0.48 mg/mL by dispersing the powder in ultrapure water (PWS 8-5 Water Distiller, Precision Water Systems). To ensure a homogeneous suspension with minimal agglomeration or precipitation, the mixture was continuously stirred for 5 minutes, followed by ultrasonication in an ultrasonic bath (Ultrasonic Cleaner, model 08896-01, Cole-Parmer) for 2 hours at 50°C.

For TiO_2_ nanoparticle mineralization, the 0.48 mg/mL suspension replaced an equivalent volume of deionized water in the mineralization solution, resulting in a final TiO_2_ concentration of 0.12 mg/mL. The system was incubated at 37°C, with samples collected at specific time points for *in situ* liquid S/TEM or *ex situ* correlative analysis. Unless otherwise stated, all reagents were obtained from Sigma Aldrich and dissolved in Milli-Q deionized water.

### Liquid-cell assembly and *ex situ* sample preparation

For *in situ* experiments, a commercially available Poseidon Select liquid holder (Protochips, Morrisville, NC, USA) was utilized. The setup incorporated a small flow management E-chip system, consisting of a Protochips ECB-55A-FM-10 bottom E-chip with a 550 × 50 μm window and 10 μm etched trenches, paired with an EPT-45W top E-chip with a 550 × 50 μm window. These E-chips were arranged perpendicularly, enhancing structural stability and reducing window bulging, albeit at the cost of limiting the viewing area to approximately 50 × 50 μm^2^.

Before use, the E-chips underwent a cleaning procedure following Protochips’ standard protocol to eliminate the photoresist protective layer. This process involved sequential immersion in acetone and methanol (each approximately 50 mL) for 2 minutes per solvent, followed by drying with compressed air. To ensure surface cleanliness and hydrophilicity, the E-chips were then subjected to plasma cleaning for 2 minutes at 30 kW using a Solarus Gatan Plasma System (Gatan, Inc., Pleasanton, CA, USA) with a gas mixture of Ne, H_2_, and Ar. This treatment promoted hydrogen bonding with water molecules, improving liquid adhesion and facilitating uniform spreading. The combination of surface tension and energy minimization ensured the formation of a stable liquid film within the system.

To assemble the liquid enclosure, 1∼2 μL of TiO_2_–CaP mineralization solution was dispensed onto the bottom E-chip and incubated for 2 minutes. Any excess liquid was removed by lightly touching the edge of a filter paper before positioning the top E-chip and sealing the liquid TEM holder. After sealing, the integrity of the E-chip window was assessed using optical microscopy (LOM). Prior to TEM imaging, the sealed liquid TEM holder underwent leak testing using a custom-designed vacuum pump station based on the HiCUBE™ Eco turbo pump platform (Pfeiffer Vacuum, Inc., Aßlar, Germany).

For *ex situ* analysis, liquid samples incubated for mineralization over time periods comparable to *in situ* observations were drop-cast onto carbon-coated 400-mesh Cu TEM grids (Electron Microscopy Sciences, Hatfield, PA, USA). These grids were pre-treated by plasma cleaning for 2 minutes at 30 kW in the Solarus Gatan Plasma System with the same Ne, H_2_, and Ar gas mixture. A 2 μL sample was deposited onto the grid and incubated for 2 minutes before being gently rinsed with methanol droplets. Afterward, the grids were air-dried and prepared for imaging.

### S/TEM Acquisition and Image Analysis

Transmission electron microscopy (TEM) and Scanning TEM (STEM) were performed using a Talos 200X TEM (Thermo Fisher Scientific, Waltham, MA, USA) equipped with four in-column SDD Super-X detectors for energy-dispersive X-ray spectroscopy (EDS) signal detection. The microscope was operated at 200 kV, with imaging and data acquisition conducted using a CETA 16M camera and Velox Software (Thermo Fisher Scientific, Waltham, MA, USA). *In situ* and *ex situ* correlative analyses were carried out using either a Poseidon Inspection single-tilt holder (Protochips, Morrisville, NC, USA) or a standard single-tilt holder (Thermo Fisher Scientific, Waltham, MA, USA).

For bright-field TEM (BF-TEM), a 100 μm objective lens was used to facilitate imaging of dry samples. Electron dosage was varied between 40-55 e/Å^2^ per acquisition. Selected Area Electron Diffraction (SAED) patterns were obtained using either a 10 μm or 40 μm aperture, corresponding to real-space diameters of approximately 125 nm and 250 nm, respectively. These SAED patterns were indexed by identifying concentric diffraction rings in polycrystalline structures, using Digital Micrograph software, and comparing interplanar spacings with reference data for TiO_2_ and HAP.

For STEM imaging, a high-angle annular dark field (HAADF) detector was utilized with a probe semi-convergence angle of 10.5 mrad. Imaging conditions included a beam current ranging from 0.02 to 0.2 nA, a pixel dwell time of 1 μs, and a typical image resolution of 1024 × 1024 pixels, resulting in a single-frame scan time of 1.3 seconds. The electron flux density (*D*) was calculated using the equation *D = It/eA*, where *I* represents the beam current, *t* is the pixel dwell time, *e* is the elementary charge, and *A* denotes the interaction area corresponding to the pixel size.^37^ Under these conditions, the electron flux density varied between 17.0 and 123 e⁻/nm^2^·s, depending on the interaction area and beam current. Videos provided as supplementary information were exported at accelerated frame rates of 10 or 50 fps. EDS maps were generated over 10 frames, with a 50 μs dwell time per pixel, and Gaussian smoothing (σ = 2; 5–7 pixel averaging) was applied to reduce noise. Background-subtracted EDS maps were automatically produced using Velox software and recorded at each tilt step. Image analysis was performed using Fiji software.^38^

## Result and Discussion

### Characterization of TiO_2_ Nanoparticles and Their Stability Under Liquid TEM

TEM micrographs (Figure 1A) revealed that after 5 minutes of continuous stirring, followed by 2 hours of ultrasonication in a 50°C ultrasonic bath to prepare the TiO_2_ nanoparticle suspension, the majority of TiO_2_ NPs in a 50 mM Tris buffer (pH 7.4) still exhibited a strong tendency to aggregate, forming clusters of varying sizes from tens to hundreds of nanometers (Figure S1). TiO_2_ is hydrophilic due to the presence of surface hydroxyl groups.^39–41^ These hydroxyl groups promote hydrogen bonding, which can drive particle-particle interactions and clumping in aqueous environments.^42^ These agglomerates consisted of assemblies ranging from a few to several hundred spherical TiO_2_ NPs (Figure 1B), with individual NPs having an average size of 17.3 ± 0.07 nm (Figure 1F). Figure 1C presents the selected area electron diffraction (SAED) pattern obtained from the TiO_2_ NPs shown in Figure 1B. The presence of well-defined concentric Scherrer rings confirms that the NPs are polycrystalline and exhibit multiple orientations, characteristic of TiO_2_. Additionally, energy-dispersive X-ray spectroscopy (EDS) mapping displays a strong Ti signal (Figure 1D and E), verifying the presence and distribution of titanium within the sample.

**Figure 1.**
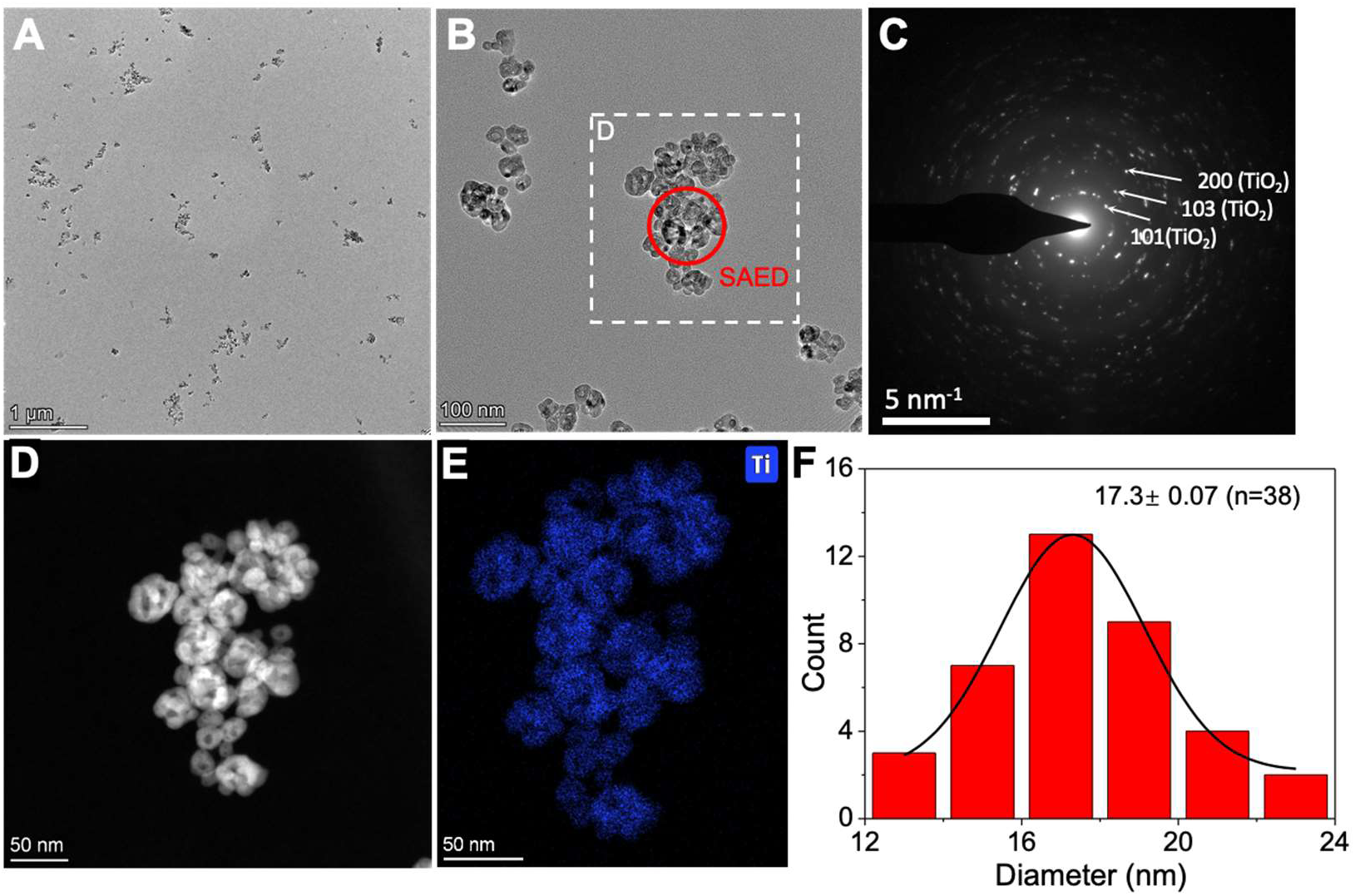
*Ex situ* characterization of pure TiO_2_ nanoparticles in 50 mM Tris-HCl (pH 7.4). (A) Representative low-magnification bright-field (BF) TEM image showing TiO_2_ nanoparticles. (B) Representative high-magnification BF-TEM image of aggregated TiO_2_ nanoparticles. (C) Selected area electron diffraction (SAED) pattern from the red-circled region in (B), confirming the crystalline nature of TiO_2_ with diffraction rings corresponding to the (101), (103), and (200) planes. (D) High-angle annular dark-field scanning transmission electron microscopy (HAADF-STEM) image of the region in (B). (E) Energy-dispersive X-ray spectroscopy (EDS) mapping, confirming the presence of Ti in the nanoparticles. (F) Particle size distribution analysis, indicating an average particle size of ∼17.3 nm.

We used liquid TEM to evaluate the stability of TiO_2_ NPs under different electron flux densities (Figure 2, Video 1 and 2) to determine optimal experimental conditions for *in situ* mineralization observations. Under low-dose electron flux density (17.7 e/nm^2^/s), TiO_2_ NPs in 50 mM Tris-HCl (pH 7.4) remained stable with minimal morphological changes after 4 minutes and 27 seconds of reaction (Figure 2A, Video 1), corresponding to an approximate cumulative electron density of 4,726 e⁻/nm^2^. In contrast, at high-dose electron flux density (123 e/nm^2^/s), the NPs rapidly expanded and moved out of the observation area within 1 minute and 40 seconds, corresponding to an approximate cumulative electron density of 18,000 e⁻/nm^2^ (Figure 2B, Video 2).

**Figure 2.**
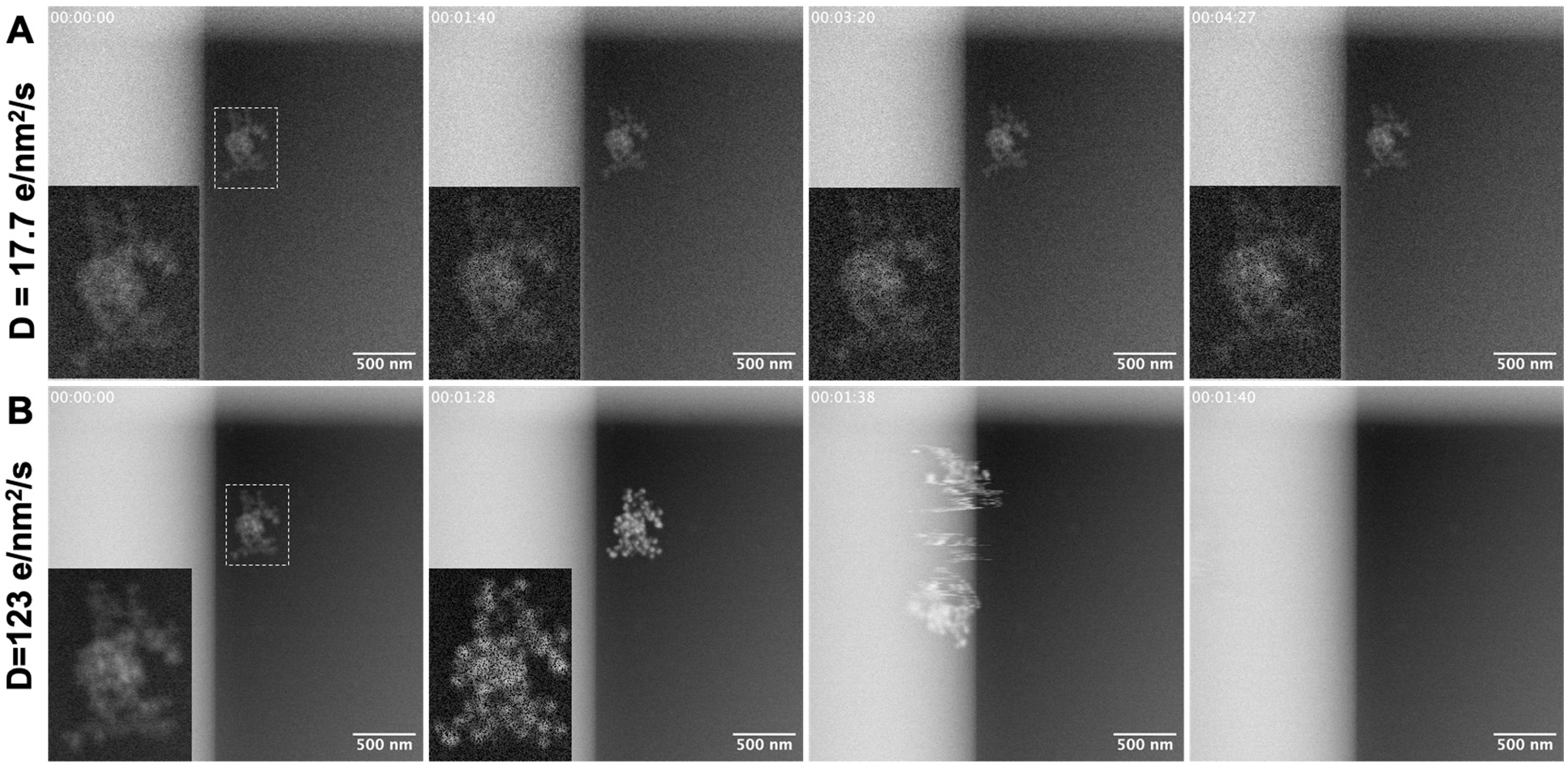
*In situ* observation of pure TiO_2_ nanoparticles in 50 mM Tris-HCl (pH 7.4) under different electron flux densities. (A) Evolution of TiO_2_ nanoparticles at a low electron flux density of **17.7 e/nm^2^/s**. Snapshots extracted from Video 1 illustrate the time-dependent structural changes at different time points. White dashed rectangles highlight regions of interest, with magnified insets providing a closer view of nanoparticle morphology over time. Under these conditions, the particles remain stable with minimal structural changes for 4 minutes and 27 seconds. (B) Evolution of TiO_2_ nanoparticles at a higher electron flux density of **123 e/nm^2^/s**. Snapshots from Video 2 reveal accelerated nanoparticle dynamics, where the particles rapidly expand and move out of the observation area within 1 minute and 40 seconds. The magnified insets offer a detailed view of the structural modifications induced by the increased electron flux.

Electron radiation damage has been shown to reduce TiO_2_ to TiO or even metallic Ti.^43,44^ Previous studies have demonstrated that electron irradiation can significantly alter TiO_2_ morphology.^45–47^ Chee. et al found, that in dry TiO_2_ sample exposed to temperatures between 500 °C to 700 °C, ∼2 Torr flowing O_2_ and only under the electron beam, 1-D TiO_2_ nanostructures formed on TiO_2_ single crystal substrates without an external Ti source.^45^ Research on the re-oxidation of reduced single-crystal TiO_2_ surfaces has revealed that the diffusion of interstitials from the bulk can drive surface morphology changes.^46,47^

In our liquid TEM imaging, the interaction between the electron beam and water generates highly reactive species that act as strong reducing agents (e.g., e⁻ₑₚ, H⁻) or oxidizing agents (e.g., •OH),^48,49^ potentially driving reduction-reoxidation processes on the surface of TiO_2_. These reactions may lead to significant morphological changes in TiO_2_ NPs. Additionally, localized heating induced by beam interactions may further accelerate these transformations, as even though bulk temperature changes are negligible, the temperature near the electron beam can rise significantly.^50–53^ Furthermore, beam interactions with water molecules generate H₃O⁺ and OH⁻, leading to charge separation and the formation of local electric fields.^48,49,54^ These fields can influence nanoparticle motion by inducing electrostatic attraction or repulsion, further contributing to instability of TiO_2_ NPs under high-dose irradiation.

### Mineralization of TiO_2_ Nanoparticles

To investigate the mineralization of TiO_2_ NPs, we performed *ex situ* TEM imaging at different time points in a CaP mineralization solution (1.7 mM CaCl_2_·2H_2_O, 9 mM K_2_HPO₄, 50 mM Tris, and 125 mM NaCl, pH=7.4) adapted from the works of Deshpande and Beniash.^36^ Early-stage imaging after a 5-minute mineralization revealed numerous aggregates ranging from 0.743 to 2.839 µm in size (Figure 3A), composed of densely packed globular particles with an average diameter of 28.16 ± 3.84 nm (Figure 3B, 3I). SAED analysis confirmed the presence of distinct diffraction rings corresponding to the (101), (103), and (200) planes, characteristic of crystalline TiO_2_ structures (Figure 3C). However, EDS mapping identified a CaP layer on the surface of TiO_2_ NPs (Figure 3D–H) that combined with the lack of crystalline peaks in SAED, indicate a layer of ACP-like phase formation on their surface.

**Figure 3.**
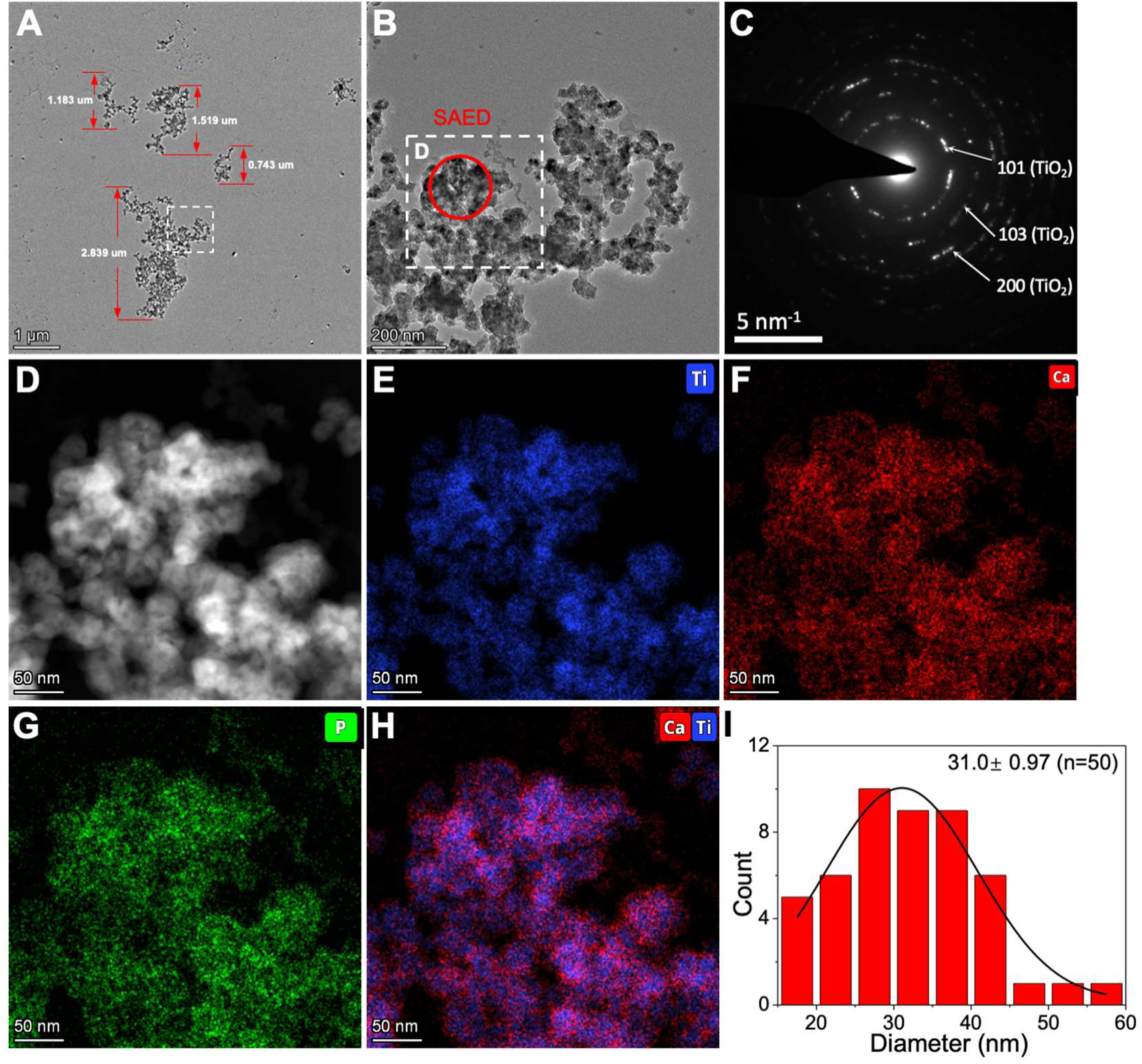
*Ex situ* observation of dispersed TiO_2_ nanoparticles after 5 minutes of reaction in a CaP mineralization solution (1.7 mM CaCl_2_·2H_2_O, 9 mM K_2_HPO₄, 50 mM Tris, and 125 mM NaCl, pH = 7.4). (A) BF-TEM image of a representative product. (B) Magnified views of the regions marked by white dashed rectangles in (A), showing packed globular particles. (C) SAED pattern from the region marked by the red circle in (B), confirming that the product remains TiO_2_ nanoparticles, as evidenced by diffraction rings corresponding to the (101), (103), and (200) crystal planes. (D) HAADF-STEM image of the region shown in (B). (E–H) EDS elemental mapping of (E) Ti (blue), (F) Ca (red), (G) P (green), and (H) an overlay of Ca and Ti, clearly highlighting a CaP layer coating the TiO_2_ nanoparticles. (I) Particle size distribution analysis, showing an average particle size of approximately 31.0 nm.

After 10 minutes, bead-like aggregates exceeding 4 µm in size were observed (Figure S2A and B). SAED analysis confirmed the presence of distinct diffraction rings corresponding to the (101), (103), and (200) planes, characteristic of crystalline TiO_2_ structures (Figure S2C), while EDS mapping again confirmed the presence of a CaP layer on the TiO_2_ surface (Figure S2D–H). Particle size analysis showed an increase in particle size to approximately 86.1 ± 1.08 nm (Figure S2I), significantly larger than the 5-minute reaction product (28.16 ± 3.84 nm, Figure 3I), suggesting continued ACP-like phase growth on the TiO_2_ NPs surface.

Extending the reaction time to 30 minutes resulted in aggregates exceeding 10 µm with a burr-like morphology (Figure 4A). The structure transitioned into needle-shaped crystals elongated along their crystallographic c-axis (Figure 4B). SAED analysis confirmed the coexistence of HAP and TiO_2_ NPs, with diffraction patterns corresponding to the (211) and (004) planes of HAP and the (101) and (200) planes of TiO_2_ (Figure 4C). EDS mapping further verified that a CaP layer enveloped the TiO_2_ NPs (Figure 4D-H). Particle size analysis revealed a continued increase in particle size to approximately 359.7 ± 25.17 nm (Figure 4I), indicating progressive mineralization and crystallization of HAP over time.

**Figure 4.**
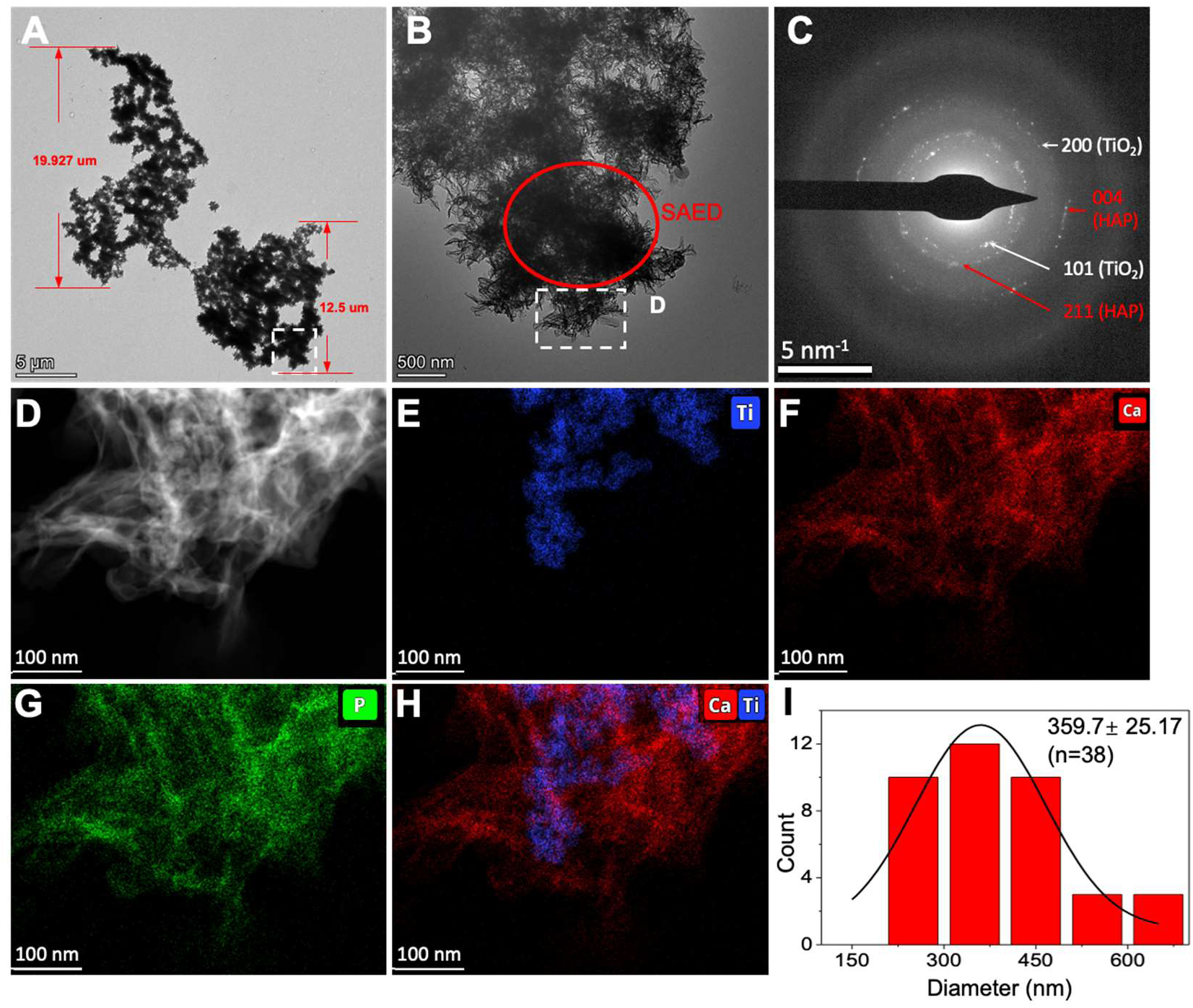
*Ex situ* observation of dispersed TiO_2_ nanoparticles after 30 minutes of reaction in a CaP mineralization solution (1.7 mM CaCl_2_·2H_2_O, 9 mM K_2_HPO₄, 50 mM Tris, and 125 mM NaCl, pH = 7.4). (A) BF-TEM image of a representative product. (B) Magnified views of the regions marked by white dashed rectangles in (A), showing a dandelion-like cluster. (C) SAED pattern from the region marked by the red circle in (B), confirming the presence of both TiO_2_ nanoparticles and HAP-like crystals, as indicated by diffraction rings corresponding to the (101) and (200) planes of TiO_2_ and the (211) and (004) planes of HAP. (D) HAADF-STEM image of the region shown in (B). (E–H) EDS elemental mapping of (E) Ti (blue), (F) Ca (red), (G) P (green), and (H) an overlay of Ca and Ti, clearly highlighting an HAP layer coating the TiO_2_ nanoparticles. (I) Particle size distribution analysis, showing an average particle size of approximately 359.7 nm.

To evaluate the role of TiO_2_ NPs in the mineralization process, we conducted a comparative *ex situ* TEM analysis of the product after 30-minute mineralization in the absence of TiO_2_ NPs (Figure S3). In this case, a large cluster structure (∼ 2 um) was observed (Figure S3A), composed of closely packed globular particles resembling a tightly clustered grapevine (Figure S3B), which remained amorphous as confirmed by SAED (Figure S3C). Additionally, EDS mapping revealed that these particles consisted primarily of CaP (Figure S3D–F). This underscores the critical role of TiO_2_ NPs in promoting HAP crystallization within the CaP system.

Correlative *in situ* liquid TEM imaging of TiO_2_ nanoparticle mineralization in the CaP solution at an electron flux density of 17.0 e⁻/nm^2^·s is presented in Figure 5 and Video 3. As previously demonstrated (Figure 2A), TiO_2_ NPs remained stable under this electron flux density (17.0 e⁻/nm^2^·s). In Figure 5, dashed lines of different colors correspond to distinct NPs or regions of interest, enabling visualization of their growth, coalescence, and movement throughout the mineralization process. At the start of the reaction, numerous nanoparticle aggregates were observed (Figure 5A). After 2 minutes and 55 seconds, these aggregates moved and coalesced into larger structures (Figure 5B). By 4 minutes and 39 seconds, aggregation 5 (black dashed outline in Figure 5) had moved out of the observation area, while aggregations 1 (red dashed outline in Figure 5C) and 3 (green dashed outline in Figure 5C) merged (Figure 5C). At 5 minutes and 38 seconds, a new aggregation (aggregation 7, dark yellow dashed outline in Figure 5D) entered the observation area and later merged with aggregation 1 at 8 minutes and 41 seconds (Figure 5D and F). Similarly, at 11 minutes and 22 seconds, another aggregation (aggregation 8, dark green dashed outline in Figure 5G) moved into the observation area and merged with aggregation 4 at 14 minutes and 19 seconds (Figure 5G and H). Between 14 minutes and 19 seconds and 36 minutes and 48 seconds, the aggregates continued to grow and became more compact over time, indicating progressive mineralization (Figure 5H and I). This corresponds to an approximate cumulative electron density of 37,500 e⁻/nm^2^ over the entire process, exceeding that of pure TiO_2_ NPs under a high-dose electron flux density (132 e⁻/nm^2^/s, Figure 2B, Video 2). This indicates that electron beam-induced effects, an intrinsic limitation of liquid-phase TEM, may also contribute to the observed phenomena. Enlarging the region of aggregation 1 (Figure S4) reveals that the NPs initially exhibited relatively smooth edges. Over time, these edges became rough and irregular, indicating the progressive phase transformation of the ACP phase into a crystalline HAP-like phase

**Figure 5.**
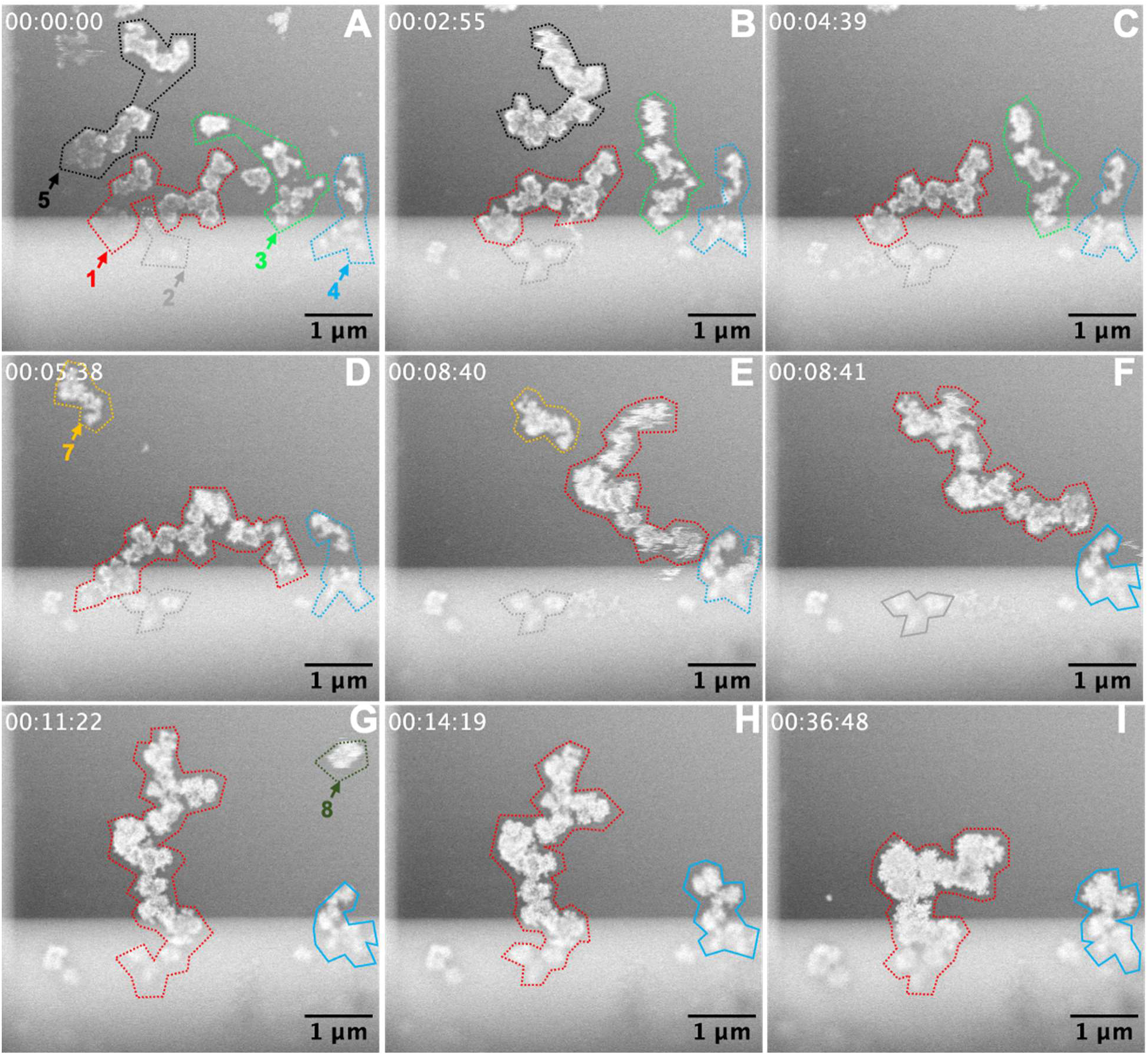
*In situ* observation of dispersed TiO_2_ nanoparticles in a CaP mineralization solution (1.7 mM CaCl_2_·2H_2_O, 9 mM K_2_HPO₄, 50 mM Tris, and 125 mM NaCl, pH=7.4) at an electron flux density of 17.0 e⁻/nm^2^·s. Time series of HAADF-STEM images captured from **Video 3** at different time points: (A) 00:00:00, (B) 00:02:55, (C) 00:04:39, (D) 00:05:38, (E) 00:08:40, (F) 00:08:41, (G) 00:11:22, (H) 00:14:19, and (I) 00:36:48. The progression highlights the morphological evolution of the nanoparticles over time. The colored dashed lines in each frame outline individual TiO_2_ nanoparticles and their structural transformations. Different colours correspond to distinct nanoparticles or regions of interest, allowing visualization of their growth, coalescence, or movement throughout the mineralization process. Black dashed outlines indicate region 5, which moves out of the observation area over time. Dark yellow and dark green dashed outlines highlight regions 7 and 8, which move into the observation area over time.

Particle size analysis revealed an initial particle size of approximately 121.6 ± 4.51 nm in the liquid sample (Figure 6A), which increased to 274.6 ± 20.28 nm after 36 minutes and 48 seconds of reaction (Figure 6B), indicating continued ACP growth and phase transformation on the surface of TiO_2_ NPs. Since the CaP mineralization solution containing TiO_2_ NPs had to be mixed before liquid-TEM observation, rather than during imaging, the initially observed particles may not be pure TiO_2_ but rather TiO_2_ particles already coated with an ACP layer. This pre-existing coating explains why the initial particle size (121.6 ± 4.51, Figure 6A) appears larger than that of pure TiO_2_ NPs (17.3± 0.07, Figure 1F). Additionally, the final particle size in the liquid sample after 36 minutes and 48 seconds (274.6 ± 20.28 nm, Figure 6B) was smaller than that in the dry sample after 30 minutes of reaction (359.7 ± 25.17 nm, Figure 4I). This discrepancy may be attributed to the confinement effects of the liquid TEM system and the limited availability of Ca^2^⁺ and PO₄³⁻ ions in the liquid sample compared to the bulk solution, which could restrict further CaP formation on the surface of TiO_2_ NPs. However, it is important to note that particle size measurements are also restricted by the limited field of view in liquid TEM, making it difficult to comment on statistical significance between samples.

**Figure 6.**
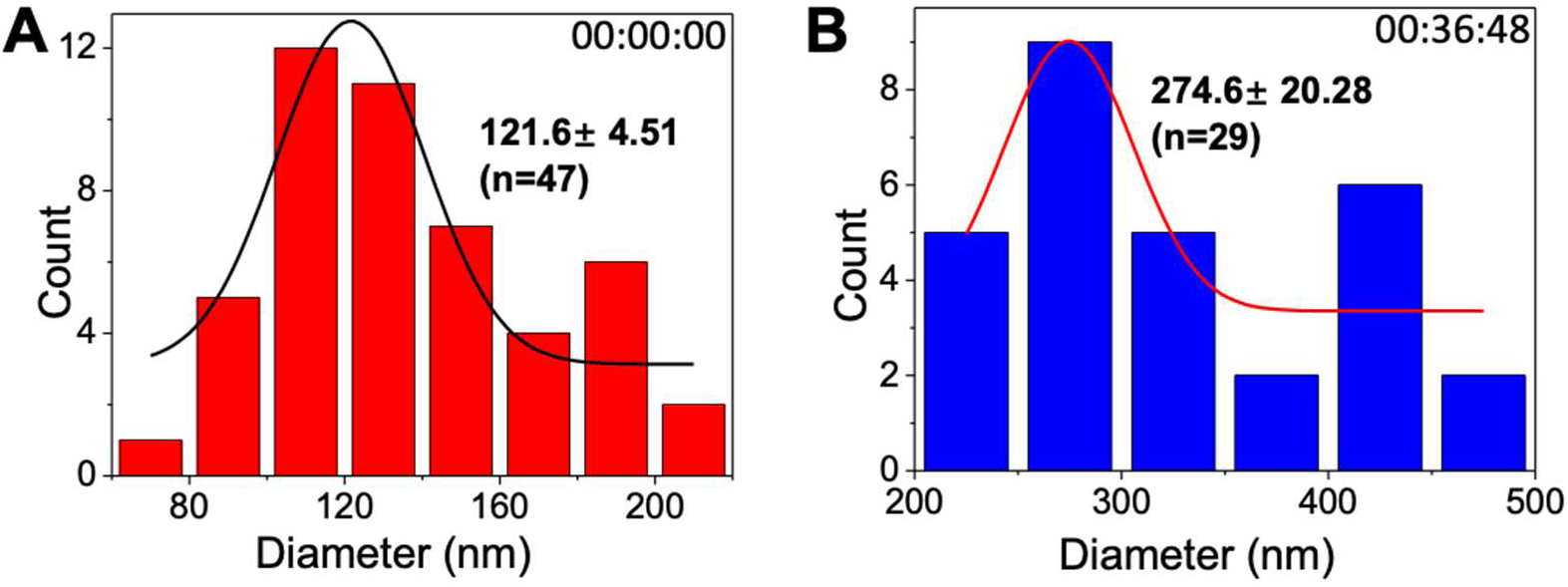
Evolution of TiO_2_ nanoparticle size distribution during *in situ* observation in Video 3 at different time points. Particle size distribution analysis at (A) 00:00:00 and (B) 00:36:48, showing an increase in average particle size from approximately 121.6 nm to 274.6 nm, respectively.

It has been reported that TiO_2_ exhibits higher surface energy than pure Ti plates, facilitating CaP nucleation on surfaces with greater surface energy.^55^ Additionally, modifications in surface characteristics, such as the increased surface area of TiO_2_ NPs compared to smooth TiO_2_ thin films, enhance CaP nucleation by providing more nucleation sites.^56^

The TiO_2_/solution interface exhibits a dipolar nature, with titanium atoms positively polarized and oxygen atoms negatively polarized. In body fluids, TiO_2_ has a strong tendency to adsorb water, leading to the formation of surface titanium hydroxide (Ti–OH) groups.^41,57^ The nature of these Ti–OH groups is pH-dependent, leading to a charge variation at the interface.^58^ The isoelectric point of TiO_2_, at which the surface exhibits zero net charge, is approximately pH 5–6. At the physiological pH of ∼7.4, the surface is slightly negatively charged due to deprotonated acidic hydroxyl groups.^41^ This negatively charged surface facilitates the adsorption of cations such as Ca^2^⁺, which may subsequently react with HPO₄^2^⁻ or H_2_PO₄⁻ to form ACP. Unlike crystalline phases, ACP possesses a disordered atomic structure, resulting in high surface energy and thermodynamic instability.^59–61^ Due to the inherent instability of ACP, it can react with free Ca^2+^ in solution, which spontaneously transforms into more stable crystalline HAP.^59–63^

## Conclusion

We propose a three-step mechanism for HAP formation on the surface of TiO_2_ NPs, as illustrated in Figure 7. This process is governed by interactions between TiO_2_ NPs and the surrounding CaP solution, ultimately leading to progressive mineralization and crystallization. Initially, TiO_2_ NPs are slightly aggregated in solution, where their high surface energy and hydrophilic nature promote interactions with Ca^2^⁺ and phosphate species. As these interactions continue, local supersaturation occurs, triggering the nucleation of ACP on the TiO_2_ surface and forming an ACP-like coated TiO_2_ complex. Over time, the inherently unstable ACP phase undergoes spontaneous transformation into crystalline HAP, driven by a reduction in surface energy. As crystallization progresses, the particles develop needle-like HAP-like structures aligned along the crystallographic c-axis. This proposed mechanism provides direct visualization of the nucleation, growth, and crystallization processes in CaP mineralization on TiO_2_ surfaces. By capturing these dynamic transformations in real time through liquid-TEM, we gain valuable insights into the physicochemical interactions governing biomineralization and bone-implant integration.

**Figure 7.**
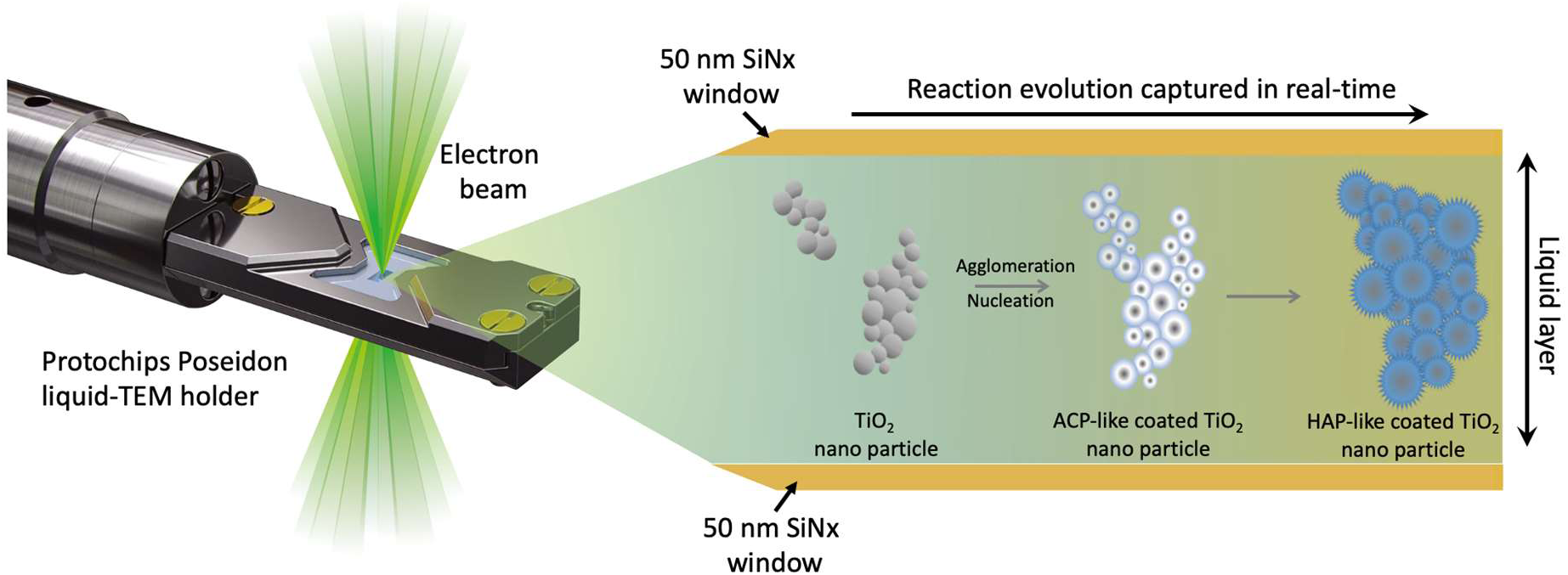
Schematic illustration of liquid-TEM observation of TiO_2_ nanoparticle evolution in a CaP mineralization solution (1.7 mM CaCl_2_·2H_2_O, 9 mM K_2_HPO₄, 50 mM Tris, and 125 mM NaCl, pH=7.4). The schematic depicts the use of a Protochips Poseidon liquid-TEM holder for *in situ* observation of the dynamic evolution of TiO_2_ nanoparticles in a CaP mineralization solution. The electron beam penetrates the liquid cell, which is enclosed by 50 nm SiNx windows, enabling real-time imaging of the reaction process. The mineralization progression is illustrated from left to right: (1) TiO_2_ nanoparticles (gray spheres) are initially aggregated in solution, (2) agglomeration and nucleation occur, forming an ACP-likelayer on the TiO_2_ surface (ACP-likecoated TiO_2_), and (3) crystallization transforms ACP into an HAP-like layer, resulting in HAP-likecoated TiO_2_ nanoparticles (blue structures). This setup allows direct visualization of nucleation, growth, and crystallization, providing insights into the mechanisms of CaP mineralization on TiO_2_ surfaces.

## ACKNOWLEDGMENTS

Financial support provided by the Natural Sciences and Engineering Research Council of Canada (NSERC) (grant no. RGPIN-2020-05722) is greatly acknowledged. Further financial support provided by NSERC for the Canada Research Chairs Program (K.G.; Tier 2 Chair in Microscopy of Biomaterials and Biointerfaces), are acknowledged. Electron microscopy was performed at the Canadian Centre for Electron.

## Author Contribution

**Conceptualization:** Jing Zhang, Liza-Anastasia DiCecco, Alyssa Williams†, Alessandra Merlo†, Kathryn Grandfield

**Methodology:** Jing Zhang, Liza-Anastasia DiCecco, Alyssa Williams†, Alessandra Merlo†

**Data curation:** Jing Zhang

**Formal analysis:** Jing Zhang, Liza-Anastasia DiCecco

**Funding acquisitions and supervision:** Kathryn Grandfield

**Writing:** Jing Zhang, Liza-Anastasia DiCecco

**Review and editing:** Jing Zhang, Liza-Anastasia DiCecco, Alyssa Williams†, Alessandra Merlo†, Kathryn Grandfield

## Notes

The authors declare no competing financial interest and no conflicts of interest.

