## Supporting Information for "Real-Time Visualization of Calcium Phosphate Formation on Titanium Dioxide Nanoparticles Using Liquid Transmission Electron Microscopy"

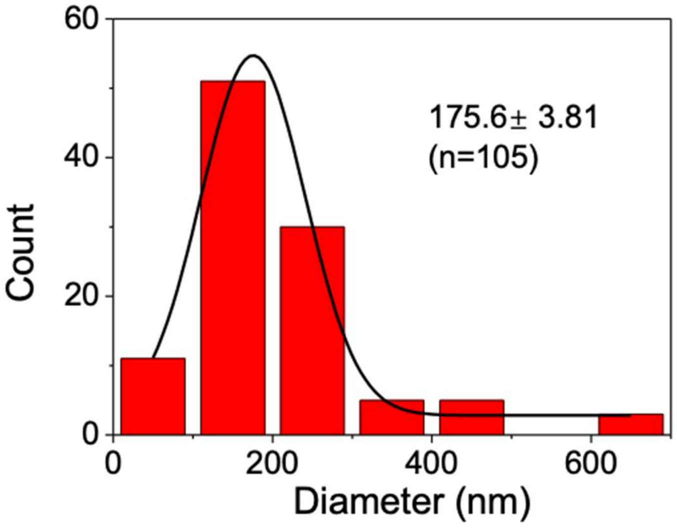


**Figure SI.** Particle size distribution analysis of $TiO_2$ nanoparticle clusters, showing an average particle size of approximately 175.6 nm.

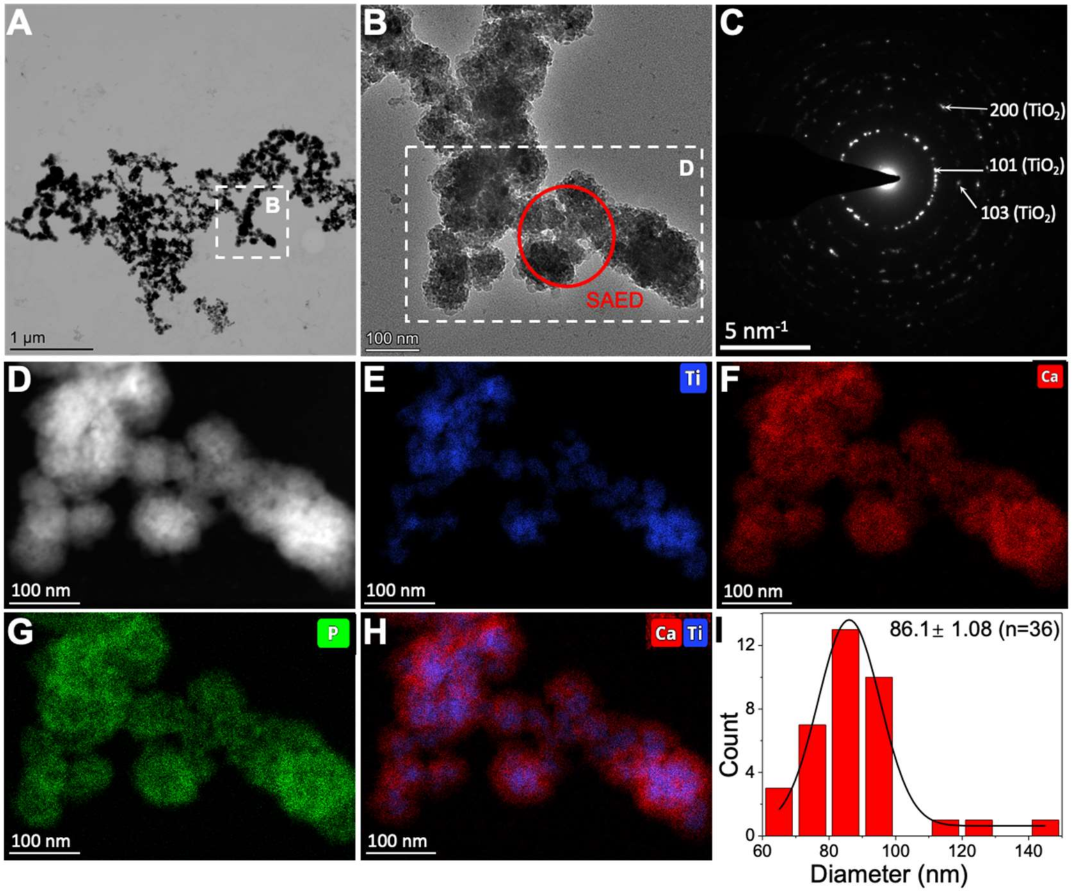


**Figure S2.** ***Ex situ*** **observation of dispersed $TiO_2$ nanoparticles after 10 minutes of reaction in a CaP mineralization solution (1.7 mM $CaCl_2 \cdot 2H_2O$, 9 mM $K_2HPO_4$, 50 mM Tris, and 125 mM NaCl, pH = 7.4).** (A) BF-TEM image of a representative product. (B) Magnified views of the regions marked by red dashed rectangles in (A), showing packed globular particles. (C) SAED pattern from the region marked by the red circle in (B), confirming that the product remains $TiO_2$ nanoparticles, as evidenced by diffraction rings corresponding to the (101), (103), and (200) crystal planes. (D) HAADF-STEM image of the region shown in (B). (E–H) EDS elemental mapping of (E) Ti (blue), (F) Ca (red), (G) P (green), and (H) an overlay of Ca and Ti, clearly highlighting a CaP layer coating the $TiO_2$ nanoparticles. (I) Particle size distribution analysis, showing an average particle size of approximately 86.1 nm.

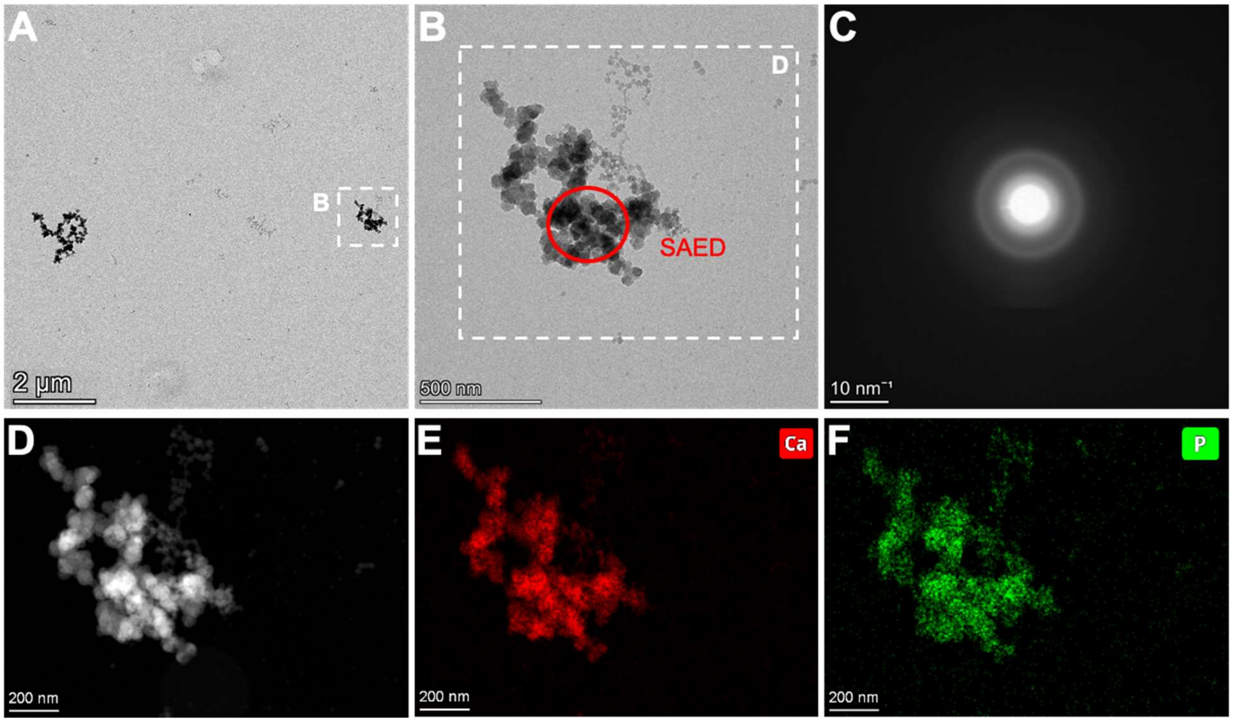


**Figure S3.** ***Ex situ*** **observation of synthesized CaP particle products after 30 minutes of reaction in a CaP mineralization solution (1.7 mM $CaCl_2 \cdot 2H_2O$, 9 mM $K_2HPO_4$, 50 mM Tris, and 125 mM NaCl, pH = 7.4).** (A) BF-TEM image of a representative product. (B) Magnified view of the region marked by the white dashed rectangle in (A). (C) SAED pattern from the region marked by the red circle in (B), indicating that the products remain amorphous. (D) HAADF-STEM image of the magnified region marked by the yellow dashed rectangle in (A). (E, F) EDS elemental mapping of (E) Ca (red) and (F) P (green), clearly highlighting that the observed particle products are composed of CaP.

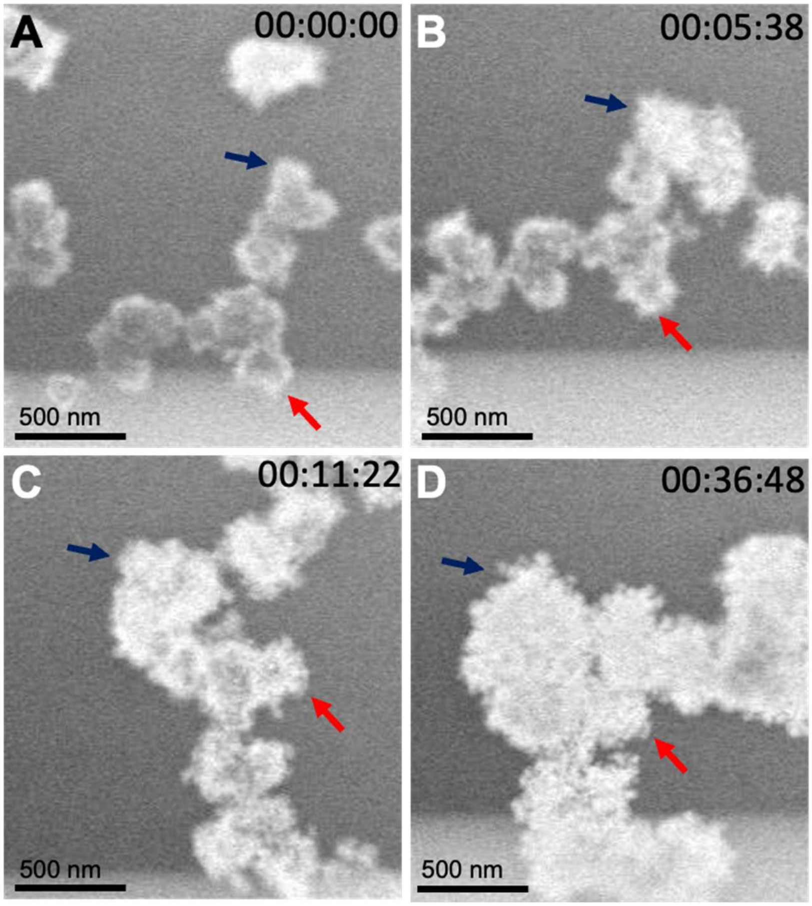


**Figure S4.** Evolution of a specific CaP-coated $TiO_2$ nanoparticle morphology during *in situ* observation in Video 3 at different time points: (A) 00:00:00, (B) 00:05:38, (C) 00:11:22, and (D) 00:36:48. Red and blue arrows highlight the location of the same particle. Initially, the nanoparticles exhibit relatively smooth edges; however, after 36 minutes and 48 seconds of reaction, they develop a rough, irregular morphology, indicating progressive ACP mineralization.

**Supporting Video Captions:**

**Video SI001**. *In situ* video of pure $TiO_2$ nanoparticles in 50 mM Tris-HCl (pH 7.4) under an electron flux density of 17.7 $e^-/nm^2/s$, displayed at 10 fps.

**Video SI002**. *In situ* video of pure $TiO_2$ nanoparticles in 50 mM Tris-HCl (pH 7.4) under an electron flux density of 123 $e^-/nm^2/s$, displayed at 10 fps.

**Video SI003**. *In situ* video of CaP-coated $TiO_2$ nanoparticles in a CaP mineralization solution (1.7 mM $CaCl_2 \cdot 2H_2O$, 9 mM $K_2HPO_4$, 50 mM Tris, and 125 mM NaCl, pH=7.4) at an electron flux density of 17.0 $e^-/nm^2 \cdot s$, displayed at 50 fps.
